# Emergence of dynamically reconfigurable hippocampal responses by learning to perform probabilistic spatial reasoning

**DOI:** 10.1101/231159

**Authors:** Ingmar Kanitscheider, Ila Fiete

**Affiliations:** Department of Neuroscience, The University of Texas, Austin, TX 78712; Department of Physics, The University of Texas, Austin, TX 78712

## Abstract

Navigation in natural environments is computationally difficult: Location errors from motion estimation noise accumulate over time, while landmarks can be spatially extended and often look alike, thus providing ambiguous data. The brain contains a number of spatially tuned neurons coding for various navigational variables, but current models do not explain how these circuits could implement navigational computations that involve non-trivial spatial reasoning. We show, using a function-first approach, that neural circuits trained to efficiently solve spatial reasoning problems with performance on par with sequential probabilistic strategies reproduce some key properties of hippocampal coding, including heterogeneous tuning, conjunctive tuning, and low-dimensional dynamics. In addition, the models predict the emergence of tuning to key latent variables that are neither present in the input data nor trained as the end result of the task, and exhibit a spontaneous dynamical reconfiguration of tuning across time during a task as the computational demands evolve, reminiscent of some of the more complex dynamics observed in the hippocampus including a switch between location and displacement coding modes. These results provide a new functional framework for understanding the rich phenomenology and potential capabilities of navigation codes in the hippocampus and associated brain areas.

## Introduction

Animals perform spatial computations to situate themselves accurately within the world, by extracting self-motion estimates from multiple senses [1], integrating these to arrive at a displacement estimate [2, 3, 4, 5, 6], then combining these displacement estimates with landmark and other data to obtain an accurate sense of their evolving location in familiar environments. These computations can be highly non-trivial when, as in the real world, the sensory data are ambiguous: Motion estimates are inherently noisy even after integrating across multiple sensory modalities [7, 8], and used on their own will result in increasingly wrong positional estimates because the integrated errors accumulate. Animals thus learn and use information about the arrangement of external landmarks to improve their location estimates [2, 9]. However, a single location can admit conflicting landmark data, and different locations – like different copses in a forest or doorways along a hall – can look alike, presenting both one-to-many and many-to-one ambiguities in spatial localization. Thus, naturalistic tasks can require the combination of multiple partially informative cues, across different points in time and space, to reason about spatial location. Algorithmically, for instance in the field of robotic SLAM [10], accurately solving such problems of agent localization with ambiguous data involves sequential probabilistic computations that continuously update multi-peaked probability distributions over the set of possible locations.

The natural behavior of animal species suggests that the brain solves these challenges efficiently: To take one example, kangaroo rats live in complicated burrow systems with multiple entrances and exits along branching dark underground passages (Fig. 1a, [11]). It is unlikely that the animal randomly explores the burrow to find an exit. Rather, despite the sensory similarities of different tunnels and various bifurcation points, the animal probably acquires and uses an internal map of the burrow, and performs spatial reasoning based on sensory data to estimate where it is and to plan how to get out.

**Figure 1:**
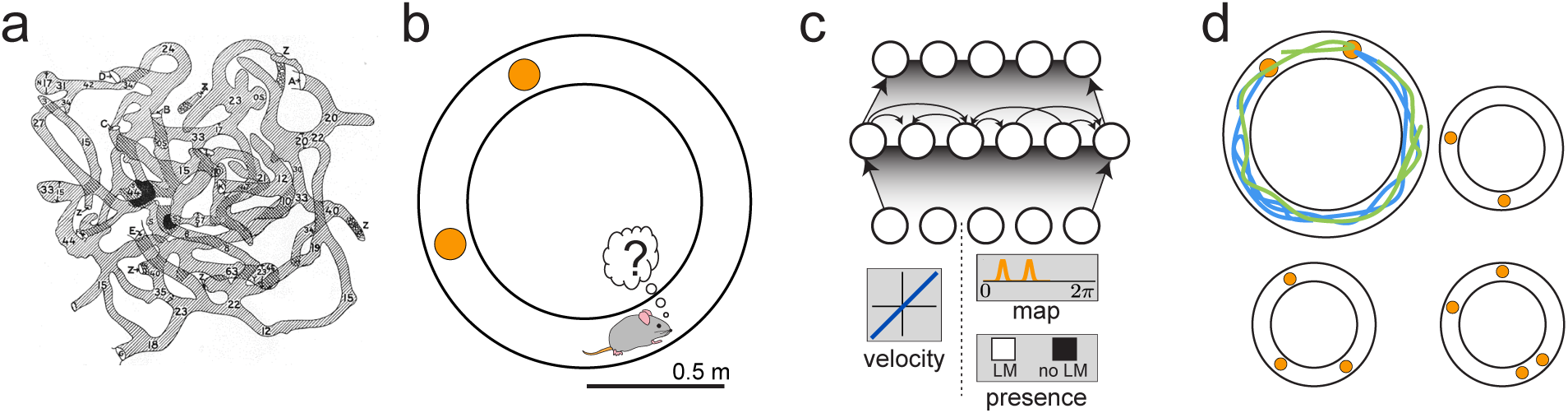
Navigation task and network model. **a**, Underground burrow of wild kangaroo rat [11]. **b**, Self-localization task: Our model animal attempts to infer its position in a known circular environment with unknown starting position, indistinguishable landmarks (orange) and noisy velocity information. **c**, Structure of the recurrent network. Input neurons encoded noisy velocity and landmark (LM) information. In the standard setup (*external map*), the LM input signaled the global configuration of LMs (*map*); in the *internal map* setup, the input only signaled whether a LM was present at the current position or not. **d**, Trajectories varied randomly and continuously in speed and direction. There were 2-4 LMs at random locations.

The brain areas involved in spatial navigation contain some of the best-characterized of cognitive codes, including head-direction cells [12, 13], place cells [14], boundary cells [15, 16], grid cells [5], border cells [17], speed cells [18] and various other cell types with spatially correlated firing patterns [19]. These cells are usually described by their tuning curves, which typically consist of bump-like activity profiles around some value of a spatial variable like head direction or location, often modulated in amplitude by signals such as animal speed or movement direction. Neural circuit models of single, homogeneously tuned populations show how, in the case of non-noisy input data, these responses may arise [20, 21, 22].

However, there are notable shortcomings in our understanding of the connection between neural tuning and the computations these circuits perform. First, tuning curves are static characterizations of neural responses, obtained by averaging over long periods of time in a fixed, known environment, and do not reveal the unspooling of the localization computation over time while non-trivial spatial inferences are being performed. The characterization of spatiotemporal activity sequences in spatial circuits [23, 24] stands apart from the static tuning-curve views of cells; while these sequences are believed to play a role in planning future paths, it is not clear that they aid in spatial localization. Second, existing models succeed in producing spatial tuning curves only if velocity inputs and corrective LM location inputs are non-noisy and sufficiently unambiguous [20, 21, 22, 25, 9, 8]. When velocity inputs are inaccurate and LM locations ambiguous, the models fail to generate localized tuning curves [8, 26]. More generally, spatial navigation involves memory, recognition, and reasoning across time within environments as well as across environments, a cognitive challenge tackled by few models (but see [27], which tackles multiple environments, although it still requires accurate sensory input). Third, tuning curves resulting from existing models are typically homogeneous, or of uniform shape. By contrast, cells in the hippocampus and associated cortical areas display many types of tuning heterogeneity: cells differ in the strength and width of location tuning [28, 29], they display conjunctive tuning to other spatially relevant variables such as head direction, velocity or environmental context [30, 31, 32] and their location tuning changes over time or with other contingencies [33, 34, 9, 35].

As a result, it is unclear how the observed cell responses underwrite localization computations in hard tasks: We lack models that perform complex spatial reasoning, to determine whether the observed phenomenology is a sufficient characterization of the responses required by the brain to solve the difficult but typical real-world problem of localization with noisy and ambiguous inputs [26].

Here we seek to understand what neural properties could allow the brain to learn, recognize and localize within different environments by training networks to localize under natural conditions of noise and ambiguity in the velocity and LM inputs. We then map the required properties to the observed phenomenology of the brain’s spatial circuits, to explore the hypothesis that a heterogeneous and dynamic neural code akin to that seen in the hippocampus arises naturally in networks that solve those functional challenges.

We set the problem of inferring location in familiar environments defined by the spatial configuration of multiple perceptually identical landmarks (LMs) along a circular track (Fig. 1b). Starting from an unknown initial location, the animal moves around the environment with imperfect motion estimates, sensing LMs that are only visible upon close approach (no long-distance vision). The animal must infer its position as it explores, in a way that is generalizable across new random trajectories and across environments with different LM configurations (Fig. 1c). We construct neurally plausible models by training recurrent neural networks to solve the problem (Fig. 1d), which, as we will show, requires integration, memorization of the spatial map (or at least the displacements between LMs), extraction of spatiotemporal context information in the form of a memory of LM encounters to disambiguate perceptually identical inputs, and multi-hypothesis probabilistic inference and reasoning. Signatures of these computations can be mapped to previously observed phenomenology in the hippocampus, providing a new functional interpretation of its neural activity in the brain’s spatial circuits.

An environment contains 2-4 identical LMs at different locations (Fig 1c). When the animal first arrives at a LM, its sensory experience is consistent with several possible locations. By continuously updating and weighting the likelihoods of these distinct possibilities, the animal could in principle resolve the ambiguity as it moved toward a second LM, based on its knowledge of the configuration and relative spacing of LMs in the environment, and of its own imperfectly estimated trajectory.

We employ a function-first approach, defining a spatial task then generating neural solutions by training recurrent networks to solve the task. The network is supplied with velocity and LM information, and asked to report on current location; neurons in the hidden recurrent network have high convergence to the output neurons, and their states are not directly constrained. The network must discover which variables to encode and how to represent and combine them to solve the task. This approach has two advantages: It generates candidate neural models for tasks where it may be difficult or impossible to hand-design a model, without imposing many constraints or assumptions, biological or otherwise, on the form the network solution might take. It also enables us to predict the encoding of emergent variables in the hidden recurrent network – key variables that are neither provided as inputs nor trained as outputs but that are necessary to solve the task, and therefore can provide a functional explanation of observed phenomenology in neural circuits.

## Results

The network receives noisy velocity inputs encoded by neurons with linear tuning curves similar to speed cells in the entorhinal cortex [18]. At a landmark (LM), the network receives positional information according to one of two schemes. In the *external map* scheme (Fig 1c schematic; used in all results except where noted), if there are *K* LMs in the environment (all assumed to be perceptually indistinguishable), then whenever the animal encounters one of the LMs, the input provides a simultaneous encoding of all *K* LM locations using spatially tuned input cells. Thus, the input encodes the *map* of the environment but does not disambiguate locations within it. This input can be thought of as originating from a distinct brain area that identifies the current environment and provides the network with its map. A trial consists of a fixed duration of exploration in a fixed environment, starting from an unknown starting location; the environment can change between trials. Environments are generated by randomly drawing a constellation of 2-4 LMs, and the network must generalizably localize in any of these environments when supplied with its map. The network must adjust its spatial inference computations on the basis of the configurations of the different environments, without changing its weights; thus, the adjustments must be dynamic.

In the *internal map* scheme (Fig 1c schematic), an input cell simply encodes by its activation whether the animal is at any LM; it does not specify the location of the LM, the identity of the environment, or the spatial configuration of the various LMs in the environment. The task in the internal map scheme is substantially harder, since the network must infer the configuration of LMs in the environment purely from the time sequence of LM visits, while simultaneously localizing itself within the environment. Information about the maps must be acquired and stored within the network. To make the task tractable, we limit training and testing in the internal map setting to four specific environments.

These two schemes allow us to test which emergent codes and computational strategies in the network are shared despite different assumptions about where spatial maps are stored. Input, recurrent and output weights are trained through supervised learning with the objective of generating outputs that correctly report location with Gaussian tuning profiles, across time (see Methods for details).

After training on random trajectories and multiple environments, the network finds a neural solution to the navigation tasks. The solution is efficient, generalizing across newly generated random trajectories and environments with a small number of neurons. Localization error, evaluated on new test trials as a function of time within the new trial, decreases relative to the unknown starting location as LMs are encountered and the network solves the location inference problem, Figure 2a (black). This performance can be compared to the much poorer performance achieved by a strategy of path integration to update a single location estimate with LM-based resets (to the coordinates of the landmark that is nearest the current path-integrated estimate), Fig. 2a, gray. The latter strategy is equivalent to existing continuous attractor integration models [27, 22] combined with a LM- or border-based resetting mechanism [36, 25, 37, 9], which to our knowledge is as far as neural models have gone in combining internal velocity-based estimates with external spatial cues. The present network goes beyond a simple resetting strategy, matching the performance of a sequential probabilistic estimator – the particle filter (PF) – which updates samples from a multi-peaked probability distribution over possible locations over time and is asymptotically Bayes-optimal (*M* = 1000 particles versus *N* =128 neurons in network; Fig. 2a, blueish-gray and reddish-gray). Notably, the network matches PF performance without using stochastic or sampling-based representations, which have been proposed as possible neural mechanisms for probabilistic computation [38, 39].

**Figure 2:**
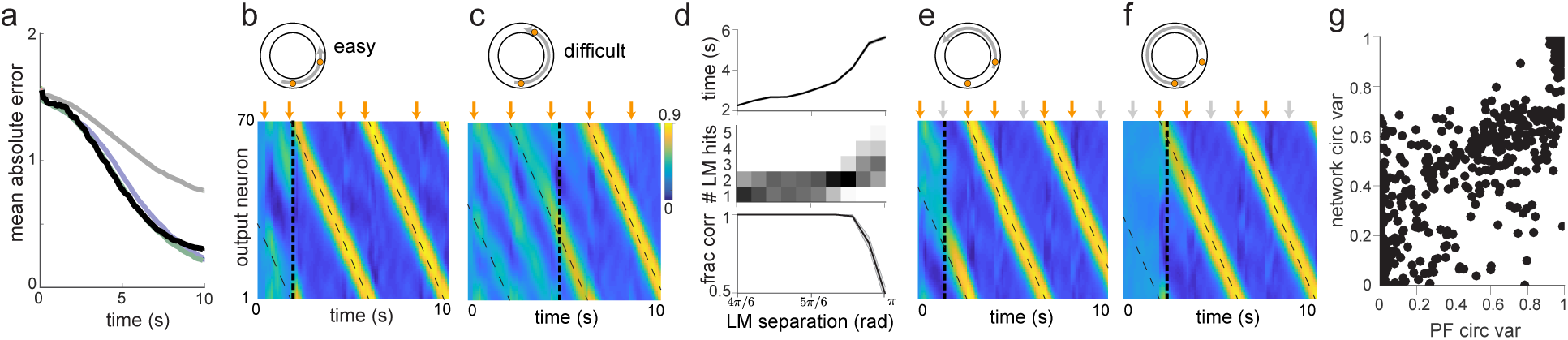
Network performance and output representation. **a**, Mean absolute localization error as a function of time within test trials, for random trajectories and LMs. Color code: network (black), path integration with reset to LM location for nearest estimated LM (gray), basic particle filter (violet), enhanced particle filter (green). **b**, Activity of output neurons ordered by preferred location as a function of time in an easy trial with two nearby LMs and a constant velocity trajectory. Orange arrows: time of LM encounters. Thick dashed line: time of disambiguation of location estimate in output layer. Thin dashed line: true location. **c**, Same as **b**, but in a difficult trial with two LMs almost opposite of each other. **d**, *Top*: Average time until location disambiguation as a function of LM separation. Standard error bars are narrower than line width. *Middle*: Distribution of the number of LM encounters until the network disambiguates location, as a function of LM separation. *Bottom*: Fraction of trials in which the network location estimate is closer to the correct than the alternative LM location at the last LM encounter, as a function of LM separation. **e**, Similar to **b**, but in a trial where the network disambiguates its location before the second LM encounter. Gray arrows mark times of LM interactions if the alternative location hypothesis had been correct. Disambiguation occurs shortly after the absence of a LM encounter at the first gray arrow. **f**, Similar to **e**, but in a trial where disambiguation occurs at the first LM location, since no LM has been encountered at the time denoted by the first gray arrow. **g**, Scatter plot of enhanced particle filter circular variance versus estimate decoded from hidden layer of the network.

To examine how the network matches the performance of a probabilistic, multi-dimensional sampling-based strategy using only relatively few deterministic units, we first study its out-puts as it localizes in a new 2-LM environment (Fig. 2b, top). Early in the trial, before any LM encounters, the output neurons exhibit no spatial tuning, Figure 2b (before 1st orange arrow). Late in the trial, a single population activity bump closely tracks the true location (Fig. 2b, after 2nd orange arrow), reflecting the network’s ability to integrate velocity cues and resolve the ambiguity associated with the identical LMs.

At intermediate times – after the first encounter with a LM – the network exhibits two simultaneous propagating bumps in the output population, corresponding to two updating hypotheses about location, starting at either of the two LM positions after the first LM encounter (Fig. 2b, between 1st and 2nd orange arrows). The network was trained to report a single location estimate, but nevertheless displays emergent multi-hypothesis coding, a signature of probabilistic computation. After the second encounter with a LM, the network somehow uses the spatiotemporal context of LM encounters in conjunction with the known map of the environment to disambiguate location, and settles on a single hypothesis by collapsing its output to a single activation bump.

Remarkably, the network’s decision on when to collapse its location estimate is flexible, and the network dynamically adapts the decision time to task difficulty: When the task is harder because of the configuration of LMs (the task becomes harder as the two LMs approach a 180 deg separation because of velocity noise and the resulting imprecision in estimating distances; the task is impossible at 180deg because of symmetry), the network keeps alive multiple hypotheses about its states across more LM encounters until it is able to reach an accurate decision, Fig. 2c (the collapse occurs after three, not two, LM encounters). Overall, the duration of multi-hypothesis representations in both time and number of LM encounters grows with task difficulty to keep accuracy high over a range of difficulties, Fig. 2d.

More remarkably, the network infers location not only by combining information from a LM encounter with its path integration estimates, but also by extracting and using information from non-encounters with LMs: If the path traversed after the first encounter with a LM corresponds to a bigger displacement than the shorter distance between two LMs, the network successfully infers that it is travelling around the far side of the track and thus disambiguates its location before the second LM encounter (Fig. 2e). If it travels for a total displacement longer than the shorter distance between the LMs before encountering even one LM, it can disambiguate its location right at the first encounter with a LM (Fig. 2f), even though the network first receives information about the configuration of LMs within the environment only at that time.

These computational capabilities go beyond those of existing hand-designed continuous attractor network models.

We next examine the dynamics of the recurrently connected hidden neurons, which enable the network’s computations. These neurons, like the outputs, implicitly compute and represent more than just a simple location estimate: Across random trajectories and environments they represent, in a way that is linearly decodable with good accuracy, the circular variance or uncertainty of the full posterior distribution of locations estimated by the PF, a second signature of probabilistic computation (*r*^2^ = 0.85; Figure 2g).

We next consider the hidden neuron states as high-dimensional *state vectors,* and project them onto a three dimensional linear subspace (using principal component analysis (PCA)) to visualize how the network structures its representations of relevant navigation variables, Figure 3. Late in a trial in an environment with two LMs (LM encounters marked by black circles), the state vector traces out a simple 1-dimensional ring that is well-parameterized by angular position on the track, showing that the network has discarded most information aside from location and the direction of movement (Figure 3a, left and Figure 3e, inset).

**Figure 3:**
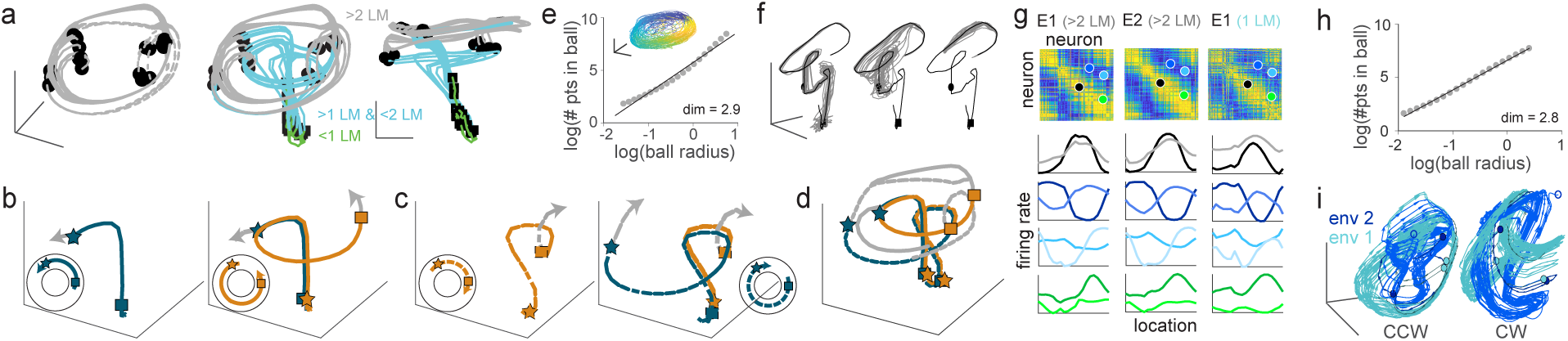
Population dynamics in hidden layer. **a**, Visualization of the full state-space dynamics of the hidden layer population, projected onto the three largest principal components, for constant-velocity trajectories. *Left*: Trajectories after second LM encounter (black circles). *Middle*: Trajectories from the beginning of the trials; black squares: first LM encounter; black circles: second LM encounter; line colors denote trajectory stage: before the first (green), between first and second (turquoise), and after the second LM encounter. *Right*: as middle, but different perspective. **b**, Divergence of trajectories for two paths that are idiothetically identical until after the second LM encounter (blue: counterclockwise travel around the short side, orange: counterclockwise travel around the long side, star and square: identities of perceptually indistinguishable LMs). Disambiguation occurs at the second encounter of the blue trajectory. See insets for geometry of trajectories and LM locations **c**, Similar as **b**, but for clockwise trajectory. **d**, All four trajectories from **b** and **c** plotted simultaneously. **e**, Correlation dimension (main plot) and state-space trajectories (inset; color corresponds to true location) after second LM encounter across environments and for random trajectories. **f**, Relaxations in state space after perturbations before the first (left), between first and second (middle) and after the second (right) LM encounter. **g**, *Top*: Correlations in spatial tuning between pairs of cells in environment E1 after the second encounter (left), in E2 after the second encounter (middle) and in E1 between first and second encounter (right). The neurons are ordered according to their preferred locations in environment E1. *Bottom*: Example tuning curve pairs (normalized amplitude) corresponding to the colored dots. **h**, Similar to **e**, but for internal map task. **i**, State space trajectories in the internal map task after the second LM encounter in two different environments. The dark green / dark blue parts of the trajectories corresponds to the sections before the third LM encounter. *Left*: Predominantly counterclockwise trajectories, *right*: Predominantly clockwise trajectories.

Yet a more complex state-space structure is visible over the full trial: states start from a common point and evolve together to an intermediate ring, moving differentially along that ring until, at the second LM encounter, they are funneled to the final 1-dimensional ring representing position along the track (Figure 3a, middle and right, trajectories colored by number of LM encounters).

The intermediate ring corresponds to times at which the output neurons represent multiple hypotheses, whereas the final location-coding ring, well-separated from the multiple hypothesis coding ring, corresponds to the period during which the output estimate has collapsed to a single hypothesis. In other words, the network internally encodes single-location hypothesis states separably from multi-location hypothesis states, but transitions smoothly between them, a novel form of encoding of probability distributions that appears distinct from previously suggested forms of probabilistic representation [38, 39].

The most interesting spatial inferences in the environment of Figure 3a occur between the first and second LM encounters, Figures 3b-d (visually indistinguishable LMs are marked differently here for expositional clarity): Consider two counterclockwise paths along very different parts of the track (Fig. 3b, blue, orange paths in insets) that are identical from the animal’s perspective (identical sensory data for motion direction and LM input) until the second LM encounter. Because of this sensory ambiguity, the state trajectories for these two paths diverge only when the blue state trajectory encounters the second LM (blue star). At this point, the location ambiguity is resolved for the blue trajectory, and the network states are funneled to the final 1-dimensional location-coding ring (gray). The orange state trajectory continues along the intermediate ring until the second LM encounter (orange square), when it is funneled onto the location coding ring (gray).

The hidden neuron state trajectories in the intermediate ring encode motion direction in a distinct way than on the final 1-dimensional location-coding ring, where the states traversed by the trajectories are nearly the same but the direction of motion is different (Figure 3a); in the intermediate ring, the states traversed in state space for clockwise and counterclockwise paths are quite separable (Figure 3b-c). Interestingly, directional information is most separable and thus readily available by the dynamically evolving representations of the hidden neuron population during this intermediate period when the network disambiguates multiple location hypotheses.

The two blue trajectories (Figures 3b-c) correspond to paths that begin at a common location on the track and also end up, at the second LM encounter, at a common location. Although the state-space trajectories start and evolve differently (Figure 3d), they converge shortly after the second LM encounter (with a slight offset because of the difference in motion direction), reflecting the fact that the network has correctly inferred the current location (although if the paths are continued forward, these trajectories will traverse the location ring in opposite directions and thus diverge again) through integration and spatial reasoning that disambiguates the LM cues.

Despite the relatively complex shape of the population state trajectories as the network solves the spatial reasoning tasks, the states occupy an overall low-dimensional manifold, as apparent in the example trajectories of Figures 3a-d. Even when summed across all environments and random trajectories, the states still occupy a very low-dimensional subspace of the full state space, quantified by the correlation dimension as *d* ≈ 3 (Figure 3e and Methods). This measure typically overestimates manifold dimension [40], and thus serves as an upper bound on the true manifold dimension. As a control, the same method yields a much larger dimension (*d* = 14) when the same network architecture is run with large random recurrent weights (Figure S6); thus, the low-dimensional dynamics is a specific, emergent property of the network when it is trained on the navigation task, and not a generic feature of networks with similar architecture. The low-dimensional state-space manifold is stable, attracting perturbed states back to it, Figure 3f, which suggests that the network dynamics follow a low-dimensional continuous attractor and the network’s computations are robust to most types of noise.

Low-dimensional population structure can alternatively be probed by pairwise neural relationships [41]: correlations or offsets in spatial tuning between cell pairs should be preserved across environments if the dynamics across environments is low-dimensional. Not surprisingly, this is the case case when we compare late-in-trial activity across environments (when the states are on the final location-coding ring in state-space), Figure 3g (first two columns: cell activities post-localization in two different environments; correlations of correlations *r* = 0.87). Surprisingly, however, the correlations for cell pairs across two different parts of a trial — the post-localization phase and the intermediate period of multiple-hypothesis representation (after the first but before the second LM encounter), when the hidden neuron states are in a different part of state space — are also well-preserved (Figure 3g, first and third columns; correlations of correlations *r* = 0.73). The computations involved over these intervals are quite different: Late in the trial, the network needs only to integrate motion cues, with simple correction of path integration noise by the already-disambiguated LMs, while early in the trial the network must disambiguate between two likely locations. The preservation of correlations across these intervals nevertheless suggests that a similar dynamical configuration of network states can perform both computations. We explore this question when examine single-cell tuning in the hidden layer.

In the *internal* map task, the network does not receive information about the location of LMs in its input; rather, it must simultaneously infer both the LM locations and the location of the animal. These determinations are inter-related, thus the much higher difficulty of the task. The higher computational complexity makes it harder to identify the same clear-cut computational stages as in the external map task, yet many state space properties are similar in the two setups. As in the external map task, the state space trajectory traces out a different low-dimensional manifold at early and late times, with different final manifolds for clockwise and counterclockwise travel (Figure S3a-d). For random trajectories, the correlation dimension *d* ≈ 3 is similar (Figure 3h), and the state-space manifold is similarly stable to perturbations (Figure S3e). As a consequence of having to code for LM locations as well, the tuning curve correlations across environments are smaller at late times (Figure S3f, first two columns; correlations of correlations *r* = 0.69) and intermediate times (Figure S3f, first and third columns; correlations of correlations *r* = 0.56) within trials.

The inference over environments can be directly visualized in state space. Similar to the external map task, in which there is a separation between hidden states for multi-hypothesis coding and hidden states coding for a single hypothesis, the network trained in the internal map task separates hidden states corresponding to different environments late in the trial, after the environments are disambiguated (Figure 3i).

To make contact with experimental phenomenology and generate predictions about the types of spatial tuning expected in systems capable of such sophisticated spatial inference, we examine the spatial tuning characteristics of the recurrently connected neurons. Because the task is to report location, we first examine how cells are tuned to location at different trial intervals: early (before the first LM encounter), intermediate (between the first and second LM encounters), and late (after the second LM encounter). Spatial tuning is dynamic, in line with changing computational demands: at early and intermediate times in the trial, a recurrently connected cell typically fires at multiple locations, seen in the broad tuning curves of Figure 4a (left, light and dark gray curves) and in the multi-modal activity distribution of Figure 4b (first column), which provides a more detailed view of spatial selectivity than the mean activity plots normally used to characterize tuning. Specifically, the activity distribution is bimodal, a reflection of the two location hypotheses that are consistent with the sensory experience of the animal. Late in the trial, as the network disambiguates the location hypotheses, the same neuron responds with single-bump tuning at a single location, Figure 4a (left, black curves) and Figure 4b (second column). Thus, the neurons are tuned to spatial location late in each trial, but not at early or intermediate times in the trial.

**Figure 4:**
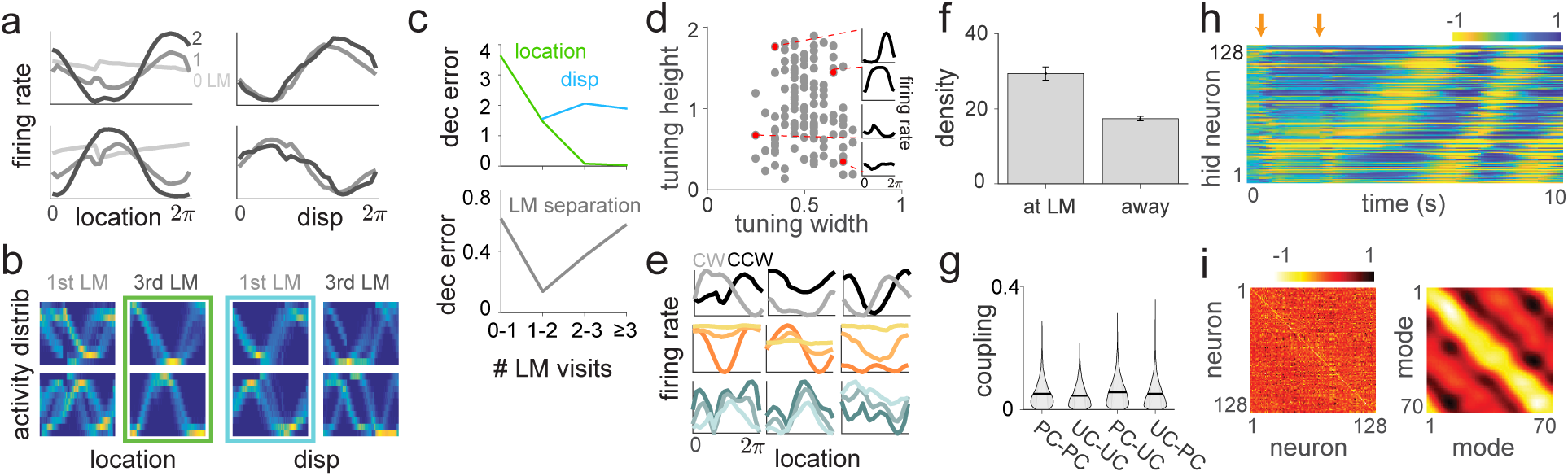
Hidden layer tuning curves. **a**, Tuning curves for location (left) and displacement from last encountered LM (right) before first LM encounter (light gray), between first and second LM encounter (dark gray), and after second LM encounter (black). The two rows correspond to two different example neurons. **b**, Distribution of activations of same neurons as in **a** conditioned on location (left two rows: after first resp. third encounter) or displacement (right two rows: after first resp. third encounter). The green and turquoise boxes mark the unimodal responses, **c**, *Top*: Square population decoding error of location (green) and displacement (blue), as a function of the number of encountered LMs. *Bottom*: Square decoding error of distance between LMs, as a function of the number of encountered LMs. **d**, Scatter plot of widths and heights of late tuning curves. Insets show example tuning curves corresponding to red dots. **e**, *Top*: Late location tuning curves for three example neurons for clockwise (gray) or counterclockwise (black) travel. *Middle*: Late location tuning curves for three example neurons for high (dark orange), middle (medium orange) or low (light orange) velocity. *Bottom*: Location tuning curves for three example neurons for high (light blue), middle (medium blue) or low (dark blue) enhanced particle filter uncertainty. **f**, Density of preferred locations close to and away from LMs in internal map task. **g**, Distribution of absolute connection strength between and across location-sensitive “place cells” (PCs) and location-insensitive “unselective cells” (UCs). The black line denotes the mean; s.e.m. is smaller than the linewidth. **h**, Hidden layer activity arranged by preferred location in an example trial. Orange arrows: first two LM encounters. **i**, *Left*: Recurrent weight matrix arranged by preferred location of neurons. *Right*: Recurrent coupling of modes defined by output weights.

Despite their poor tuning to location at intermediate times, the same two neurons respond with clear, single-bump tuning to a different variable in this interval: The displacement from the last encountered LM, Figure 4a (right, dark gray curves) and Figure 4b (third column). By contrast, at late times, when displacement from the last encountered LM ceases to be an important variable for spatial inference, the tuning to displacement becomes bimodal, Figure 4b (fourth column). The changing importance of different variables at different times, and the dynamical encoding of these variables by the network, can also be observed and quantified in a different way: By linearly decoding the relevant variables from the neural population. The squared error in a linear decoding of location decreases with LM visits, while the error in estimating displacement initially decreases then levels off, Figure 4c (top). In sum, the network switches from a displacement coding strategy at early times to encoding location at later times in each trial.

Displacement from the last encountered LM is supplied as neither an input to the network nor enforced by training as a desired output. However, it is a key variable required to solve the ambiguous LM problem, when combined with information about heading direction, which we noted earlier (Figure 3b-c) is also most linearly decodable during the intermediate period in each trial. To see why, consider again the environment shown in the insets of Figure 3b-d, with identical LMs near 11 and 3 o’clock. If, *traveling counterclockwise* after hitting a LM (either trajectory in the two insets of Figure 3b), the animal traverses a *short distance* before hitting another LM, the present location is 11 o’clock (Figure 3b, left inset trajectory). On the other hand, a *long distance* to the second LM encounter would imply that the present location is 3 o’clock (Figure 3b, right inset trajectory). The network discovers that displacement from the last LM encounter is a key latent variable, and its encoding is an emergent property. Intriguingly, a similar displacement-to-location coding switch has been observed in [42], suggesting that the empirically observed switch may be related to the brain performing spatial reasoning to disambiguate between multiple location hypotheses.

Another key latent variable, not supplied as an explicit input to the network, is the spacing between LMs in a given environment. The network must compare its estimate of displacement between consecutive LM encounters with the spacing of LMs to disambiguate location. In environments with two LMs, we find that LM spacing is computed based on the map supplied to the network on each trial, and encoded in a way that can be readout by a linear decoder that remains fixed across trials and environments. The representation is particularly accurate around the time just before and after the first LM encounter, when location disambiguation takes place, Figure 4c (bottom), similar to the dynamically modulated decodability of displacement (Figure 4c, top). In sum, the tuning to key features is highly reconfigurable over time in a task, as the computational demands shift, even though the state-space structure is low-dimensional at any given time.

Unlike hand-designed continuous attractor networks, in which neurons typically display homogeneous tuning across cells [20, 21, 22], the present model naturally reproduces the many types of heterogeneity observed in hippocampus and associated cortical areas. Network tuning curves exhibit a wide distribution of widths and heights (Figure 4d), and many hidden cells show conjunctive tuning to location, direction, and velocity (Figure 4e, top and middle rows, respectively and Figure S5), despite the overall low-dimensional dynamics of the network (Figure 3e). In particular, the direction selectivity of hidden units mimics the direction selectivity of place cells in 1D tasks [30]. By contrast, a network trained on 2D spatial tasks shows weak or no direction selectivity in location tuning [43], similar to the lack of direction selectivity of place cells in 2D open fields [44]. The hidden units also show conjunctive tuning to uncertainty, with uncertainty defined by the circular variance of a particle filter run on the same trajectory and LM configuration (Figure 4e, bottom row).

An analysis of the distribution of recurrent weights shows that groups of neurons with strong or weak location tuning or selectivity have similar patterns and strengths of connectivity within and between groups, Figure 4g, again in contrast to what one would expect to obtain from hand-designed attractor network models. However, the result is consistent with data suggesting that place cells and non-place cells do not form distinct sub-networks, but are part of a system that collectively encodes more than just place information [45].

Comparing location tuning and recurrent connectivity in the hidden layer reveals another aspect of its mixed representation. Late in trials, if the recurrently connected neurons are ordered according to their preferred locations, they exhibit an activity bump that moves coherently with the network’s location estimate (Figure 4h). However, the recurrent weights between these neurons do not exhibit the characteristic circulant matrix structure that would underlie a travelling activity bump in hand-designed continuous attractor models, Figure 4i (left). A circulant matrix structure does exist, but it is shuffled by mixture components defined by the output weights, Figure 4i (right): Connections between appropriate neural mixtures in the hidden layer – defined by the output projection of the neurons – exhibit a circulant structure, but the actual recurrent weights do not, even after sorting neurons according to their preferred locations. Thus, the network implements a generalization of hand-wired attractor networks, in which the integration of velocity inputs by the recurrent weights occurs in a basis shuffled by an arbitrary linear transformation. Given these results, one cannot expect a connectomic reconstruction of a recurrent circuit to display an ordered matrix structure even when the dynamics are low-dimensional, without taking into account the output projection.

Hippocampal studies have shown that place cells accumulate near certain locations in an environment, such as at reward sites [46, 47, 48]. Less established is the question of whether place fields are assigned at a higher rate to locations of more general strategic importance on a task. We find that the density of preferred locations of hidden layer neurons in both the external and the internal map networks is significantly higher close to LM positions than away from them (Figures 4f and S1f). Thus, we expect a statistically higher density of place fields at strategic or spatially informative locations, regardless of whether these locations are intrinsically rewarding, across hippocampal areas involved in storing and representing spatial maps, as well as in areas that perform spatial inference given the map.

Aside from differences in the propensity of the internal map network to encode and store environment-specific information about LM locations, the external and internal map settings result in similar emergent codes and dynamics, including the transition from displacement-to-location coding (Figure S1a-c), and an increase in population-level information about LM decoded locations (Figure S1c, bottom), with increasing numbers of LM encounters within a trial. Like the external map network, the internal map network also shows high degrees of heterogeneous (Figure S1d) and conjunctive (Figure S1e) tuning, clustering of preferred locations of hidden layer neurons close to LM positions (Figure S1f) and similar patterns of connectivity between strongly and weakly location-tuned cells (Figure S1g).

Another way to probe the generality of the network predictions is to change the neural nonlinearity or impose other constraints on neural activity. For the external map setup, we restricted the activity of neurons in the hidden layer to be non-negative, similar to firing rate in biological neurons, to obtain similar results with respect to location and displacement tuning (Figure S2a-b), the transition in linear decodability of displacement to location from the population (Figure S2c, top), the dynamically varying decodability of LM separations within trials (Figure S2c, bottom), the presence of heterogeneous (Figure S2d) and conjunctive tuning (Figure S2e), lack of modularity in connectivity between cells with high and low amounts of spatial selectivity (Figure S2f), and the preservation of cell-to-cell correlations across time within trials and across environments (Figure S2g). The nonlinearity does affect the distribution of recurrent weights: The distribution of non-diagonal elements in the non-negative network is sparse (excess kurtosis *k* = 7.8), while it is close to Gaussian for the external and internal map networks with *tanh*-nonlinearity (*k* = 0.6 and *k* = 0.9 respectively; Figure S4a); however, the distributions of eigenvalues of the recurrent weights have similar characteristics for all trained networks (Figure S4b).

## Discussion

We have studied the emergence of neural phenomenology from a function-centric perspective, by training neural networks to recognize and localize efficiently within ambiguous environments using impoverished and noisy sensors. Our study is inspired by past work on how neural-like responses can result when networks are trained on vision [49, 50, 51] or decision-making [52] tasks. However, our focus has been on cognitive areas rather than sensory ones, and on computations that involve memory (and thus a recurrent dynamics) and probabilistic inference.

Our results transcend previous modeling approaches that implement path integration with boundary correction using continuous attractor networks [27, 22, 9] in several ways: First, our model efficiently deals with noise and ambiguity, reaching similar performance as probabilistic, sampling-based strategies. Second, the process of network training finds a generic solution subject to the specified the computational constraints and naturally reproduces many aspects of tuning curve heterogeneity and conjunctive tuning observed in hippocampus and associated areas, while generating predictions about time-resolved phenomena like the reshaping of neural responses over the course of a task based on task demands. Third, the result shows that measuring tuning curves is by itself unlikely sufficient to identify the computational strategy a neural area implements [26], and that a population decoding approach can reveal more about the encoded variables and how they evolve over time.

It is reasonable to wonder whether the emergent coding observed in the hidden layer depends in detail on the network architecture including the single-neuron model and the types of inputs the networks receives and how they are encoded. To address this concern, we trained networks to solve the multi-environment localization problem using different inputs (internal versus external map tasks) and constraints (unconstrained versus non-negative activation in neurons). The predictions for hidden layer tuning and population dynamics were remarkably consistent, with slight differences explainable by differences in the computational task associated to each architecture.

The networks presented here do not incorporate certain biological constraints, such as spiking neurons or Dale’s law, but nevertheless reproduce a number of key properties of hippocampal coding. This suggests that the emergent coding of spatial variables in the brain may be shaped in significant part by the computational demands of navigational inference rather than detailed biological constraints.

## Methods Summary

An animal runs with varying velocity in a circular environment starting from a random, unknown position and eventually infers its position using noisy velocity information and two, three or four indistinguishable LMs. In the *external map* task, LM locations were random and the set of locations (map) were provided to the network, whereas in the *internal map* task one of four LM configurations was used, but the maps were not provided to the network. LMs could only be observed a short distance. A three-layer network with a recurrent hidden layer was trained to infer location. Velocity and LM encounter information were encoded in the input layer, and all weights of the network were trained. The training target for the output layer was activation of a unit with von Mises tuning and preferred location matching the true location.

Network performance was compared to a number of alternative algorithms: *Path integration + correction* integrated the noisy velocity information starting from an initial location guess and corrected this estimate by a reset to the coordinates of the nearest LM when a LM was encountered. *Particle filters* approximated sequential Bayesian inference given the available velocity and LM information, with each particle capturing a location hypothesis whose posterior probability is given by an associated weight. Particle locations are updated using velocity information and particles are reweighted after LM encounters. The *enhanced particle filter* also reweights particles when a LM is expected but not encountered, thus can infer location not only from the presence but also from the absence of LMs.

The output and hidden representations of the trained network were evaluated in a variety of conditions involving both random and fixed LM locations and trajectories with random and fixed velocities.

## Methods

### 1 Definition of environments and trajectories

The task is defined by a simulated animal moving along a circular track of radius 0.5 m for 10 seconds. The animal starts at a random, unknown position along the circle at rest and starts running along a trajectory at non-constant velocity. A trajectory is sampled every *dt* = 0.1s in the following way: At each time *t*, acceleration *a_t_* is sampled from a zero-mean Gaussian distribution with standard deviation 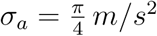 that is truncated if 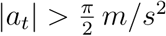. Acceleration is integrated to obtain the velocity *υ_t_* and truncated if 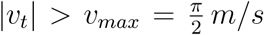. The actual location on the track is the integral of this velocity.

In a trial of the *external map* task, the locations of *K* = 2, 3, or 4 indistinguishable LMs were determined sequentially: the first LM was sampled from a uniform random distribution on the circle, with subsequent LMs also sampled from a uniform random distribution but subject to the condition that the minimum angular distance from any previously sampled LM is at least *δ* = *π*/9 rad.

The *internal map* task involved four environments, each with a unique configuration of LMs: two environments had two LMs, one had three and the last had four. LM locations in the four environments were chosen so that pairwise angular distances were sufficiently unique to allow the inference of environment identity. LM coordinates in environment *e_i_* were given by: *e*_1_ = {0, 2*π*/3} rad, *e*_2_ = {1.9562, 3.7471} rad, *e*_3_ = {0.2641,1.2920, 3.7243} rad, *e*_4_ = {3.0511, 3.8347, 5.1625, 5.7165} rad.

### 2 LM observation

The animal is considered to have encountered a LM if it approached within *d_min_* = *υ_max_* · *dt*/2 = *π*/40 *m*/2 = *π*/20 rad. This threshold is large enough to prevent an animal from “missing” a LM even if it is running at maximum velocity. Hovering around the same LM or approaching the same LM consecutively would only trigger a LM encounter at the first approach; a new encounter was only triggered if the animal approached a LM different than the previous one. Also, only trials in which the animal encountered at least two different LMs were included.

### 3 Sensory noise

The largest sources of uncertainty in the tasks were the unknown starting position and the indistinguishability of the LMs. In addition, we assumed that the velocity information and the LM-location memory (in the external map scenario) were corrupted by noise. At each time step of size *dt* = 0.1, the velocity input to the network corresponded to the true displacement *vdt* corrupted by zero-mean Gaussian noise of standard deviation *σ_υ_* = *υ_max_dt*/10. In the external map task, the LM map provided to the network and particle filter was corrupted by zero-mean Gaussian noise with standard deviation *σ_l_* = *π*/50 rad, without changing the relative LM positions: The map was coherently slightly rotated at a LM encounter, and the rotation was independently sampled at each LM encounter.

### 4 Network architecture and training

The network consisted of three layers of rate neurons with input-to-hidden, hidden-to-hidden and hidden-to-output weights. All weights were trained.

#### Network input

The input layer consisted of 80 neurons in the external map case and 11 neurons in the internal map case. Ten neurons coded for velocity corrupted by noise (noise as described above). The velocity neurons had a minimum firing rate between 0 and .2 and a maximum firing rate between .8 and 1 in arbitrary units, and within this output range coded linearly for the whole range of velocity between –*υ_max_* and *υ_max_*. Negative and positive velocity here corresponds to clockwise and counterclockwise travel respectively.

The remaining neurons (70 in the external map case and 1 in the internal map case) coded for LM input and were activated only at the time step of and up to three time steps after a LM encounter.

In the external map case, the LM input simultaneously encoded the locations of all LMs in the environment, thus supplying a map of the environment, but contained no information about which LM was currently encountered. The LM neurons had von Mises tuning with preferred locations *x_j_* = (*j* — 1) · 2*π*/70 rad, *j* = 1…70, that tiled the circle equally. Given *n* LMs at locations *l_i_ i* = 1…*n*, the firing rate of the *j*-th LM input neuron was given by

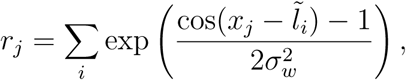

where 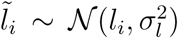 is the noise-corrupted LM coordinate (see “Sensory noise” section). This mixture of von Mises activation hills produces the pattern depicted as the “map” input in Figure 1c.

In the internal map case, the LM input neuron consisted of a single binary neuron that responded for four time steps with activation 1 in arbitrary units whenever a LM was encountered. This input encoded neither environment identity nor LM location.

#### Hidden layer

The hidden layer consisted of 128 recurrently connected neurons. The activation **h**_*t*_ of hidden layer neurons at time step *t* was determined by

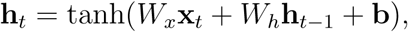

where **x**_*t*_ are the activations of input neurons at time step *t*, *W_x_* are the input-to-hidden weights, *W_h_* are the hidden-to-hidden weights and **b** are the biases of hidden neurons. The nonlinearity should be considered as an effective nonlinearity at long times; since the time step *dt* = 0.1s was large compared to a typical membrane time constant (*τ* ≈ 0.02s), we did not include an explicit leak term.

#### Hidden layer (non-negative network)

In the non-negative network (Figure S2), the recurrent activation was determined by

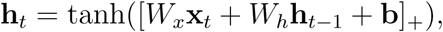

where [*u*]_+_ denotes rectification.

#### Output layer

The output layer consisted of a population of 70 neurons with activity **o**_*t*_ given by

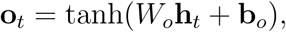

where *W_o_* are the output weights and **b**_*o*_ the biases of the output neurons.

#### Network training

The training targets of the output layer were place cells with von Mises tuning of width *σ_o_* = *π*/6 rad to the true location *y_t_*,

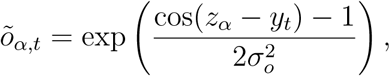

where *z_α_*, *α* = 1…70 are the equally spaced preferred locations of each training target.

The network was trained by stochastic gradient descent using the Adam algorithm [53], to minimize the average square error between output **o**_*t*_ and training targets 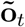, with the average taken over neurons, time within each trial, and trials. The gradients were clipped to 100. The training set consisted of 10^6^ independently generated trials. During training, performance was monitored on a validation set of 1000 independent trials and network parameters with the smallest validation error were selected. All results were cross-validated on a separate set of test trials to ensure that the network generalized across new random trajectories and/or LM configuratio/sans.

#### Network location estimate

Given the activity of the output layer at time *t*, we define the network location estimate for that time to equal the preferred location (the preferred location was set over training) of the most active output neuron:

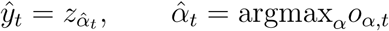

### 5 Performance comparisons

In figure 2a, we compared the performance of the network in the external map task with a number of alternative algorithms. To ensure a fair comparison, we make sure that each alternative algorithm has access to exactly the same information as the network: the LM identities are indistinguishable and both velocity and LM location information are corrupted by the same small amount of sensory noise.

#### Path integration + correction

This algorithm implements path integration and LM correction using a single location estimate, similar to what is implemented in hand-designed continuous attractor networks that implement resets at boundaries or other LMs [37, 36, 25, 9]. The algorithm starts with an initial location estimate at *y* = 0 (despite the true initial location being random and unknown), and integrates the noise-corrupted velocity signal to obtain location. At each LM encounter the algorithm corrects its location estimate to equal the coordinates of the LM nearest to its current estimate.

#### Basic Particle filter

Particle filters implement approximate sequential Bayesian inference using a sampling-based representation of the posterior distribution. Here, the posterior distribution over location at each time point is represented using a cloud of weighted particles, each of which encodes through its weights a belief, or estimated probability, of being at a certain location. In the beginning of the trial, *N_p_* = 1000 particles are sampled from a uniform distribution along the circle and weighted equally. In the prediction step, particles are independently propagated using a random walk whose mean is the noise-corrupted velocity update and whose standard deviation is the velocity noise *σ_υ_*. In the absence of a LM encounter, particle weights remain unchanged and the particle cloud diffuses. If a LM is encountered, the importance weights *ω_t,β_* of particles *β* =1…*N_p_* are multiplied by

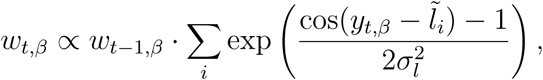

where *y_t,β_* are the current estimates of the particles, and the weights are subsequently normalized such that *∑_β_ ω_t,β_* = 1. If the effective number of particles becomes too small, i.e. 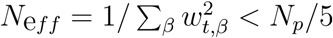 the particles are resampled using low variance sampling [10] and the weights equalized. This resampling step both allows for better coverage of probabilities and permits the particle cloud to sharpen again. The particle filter estimate at a given time point is given by the weighted circular mean 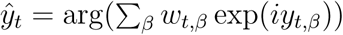 of the particle locations. In addition we also calculate the circular variance using 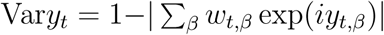.

#### Enhanced particle filter

This particle filter has identical initialization, prediction step and weight update at LM encounters as the basic particle filter and proceeds in exactly the same way until the first LM encounter. Subsequently, the enhanced particle filter can also use the absence of expected LM encounters to narrow down its location posterior, similar to the network’s ability shown in Figure 2e-f. This is implemented in the follow way: If a particle comes within the observation threshold *δ* of a possible LM location but no LM encounter occurs, the particle is deleted by setting its weight to zero; afterwards the particle weights are renormalized. A complication to this implementation is that a subsequent LM encounter only occurs if the current LM is different than the previous one (see section “Landmark encounters”); to prevent the deletion of particles that correctly report a LM at the current position but do not receive a LM encounter signal because it is the same LM as previously encountered, particles are only deleted if they come within the observation threshold δ to a possible LM that is different than the last LM and do not encounter it.

In case all particles have been deleted, particles are resampled from a uniform distribution and their weights are equalized. As for the basic particle filter, particles are resampled whenever the effective number of particles becomes too small, 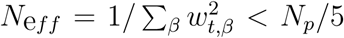. Also the particle filter estimate 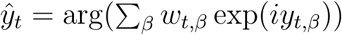 and the circular variance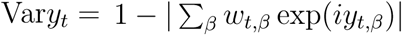 is calculated in an identical way.

### 6 Analysis of location disambiguation in output layer

The timing and accuracy of location disambiguation in Figure 2 was calculated in the following way. We first constructed the trajectory of the “alternative location hypothesis”, corresponding to the location estimates of a model animal that made the wrong location disambiguation at the first LM encounter, but otherwise updated its location by the correct velocity. This trajectory is shifted relative to the true trajectory by a constant distance equal to the distance between the two LMs. At each point in time, we then identified the two neurons in the output population whose preferred locations were closest to that of the true and alternative trajectory, respectively; the activation of these neurons roughly corresponded to the height of the activation bump corresponding to the true and alternative location hypothesis as seen in Figure 2b and c. The disambiguation time was defined as the earliest time after which either the true or alternative location bump height fell below a threshold of 0.1 and stayed beyond that threshold until the end of the trial.

To determine the accuracy of location disambiguation the network estimate at the last LM interaction was analyzed. If this network estimate was closer to the true than to the wrong LM location the trial was categorized as a correct trial, otherwise it was categorized as an incorrect trial.

### 7 State space analysis

We performed principal components analysis (PCA) on the hidden neuron states from training trials to obtain the top three principal directions. We then projected network states obtained from the distribution of testing trials 2 or 3 (see SI) onto these principal directions. The resulting reduced-dimension versions of the hidden neuron states from testing trials are shown in Figure 3 and S3.

### 8 Correlation dimension

To calculate the correlation dimension we first performed linear dimensionality reduction (PCA) on hidden layer activations from the training trials, retaining 20 principal components. In the 20-dimensional space, we randomly picked 1000 base points. From each of these base points, we estimated how the number of neighbors in a ball of radius *R* scales with *R*. The minimum ball radius was determined such that the logarithm of the number of neighbors averaged over base points was near 1. The maximum radius was set to 10 times the minimum radius, and intermediate values for the radius were equally spaced on a log-scale. The slope of the linear part of the relationship between the logarithm of number of neighbors versus ball radius determined the fractal dimension.

## Supplementary Information

### Overview over testing distributions

After training, the networks were evaluated in different testing configurations that each consisted of a distribution over landmark configurations and trajectories:

1. Training distribution: This test set was generated exactly as in the training set, as described in section “Definition of environments and trajectories”.
2. Fixed landmarks, random trajectories: The landmark configuration was given by two landmarks located at *e* = {0, 2*π*/3}, the trajectories were sampled in an identical way as in the training distribution. Note that this landmark configuration corresponds to the first environment in the internal map task.
3. Fixed landmarks, constant velocity trajectories: The landmark configuration was given by two landmarks located at *e* = {0, 2*π*/3} and the trajectories were given by constant velocity trajectories with |*υ_t_*| = *υ_max_*/2. The initial position and the direction of the trajectory was random.
4. Two variable landmarks, constant velocity trajectory: The landmark configuration was given by two landmarks located at *e* = {0, 2*π*/3 + *απ*/3}, where α ∊ [0,1]. The trajectories were given by constant velocity trajectories with |*υ_t_*| = *υ_max_*/2 and the initial position and the direction of the trajectory was random.
5. Two environments, random trajectories: The landmark configuration was given by either *e*_1_ or *e*_2_ of the internal map task, trajectories are random

### Simulation parameters of figures

- *Figure 2a*: 5000 trials sampled from training distribution (Distribution 1).
- *Figure 2 b,c,e,f*: Example trials from distribution 4 with different values of landmark separation parametrized by *α*.
- *Figure 2d*: 10000 trials from distribution 4, 1000 for each of the 10 equally spaced values of *α*.
- *Figure 2g*: 4000 trials from distribution 1 were used to train a linear decoder to predict the posterior circular variance of the enhanced particle filter from the activity of the hidden units. The performance of the decoder was evaluated on 1000 test trials. The output of the linear decoder was clipped to be between 0 and 1 to conform to the range of the circular variance.
- *Figure 3a-d*: 1000 trials from distribution 3; the sensory noise was set to zero. In Figure 3b-d, the starting point was chosen such that the simulated rat would travel a fixed distance before encountering the first landmark.
- *Figure 3e,h*: 5000 trials from the distribution 1.
- *Figure 3e, inset*: 100 trials from the distribution 1.
- *Figure 3f*: For the base trial, a trial with two landmarks and random trajectory was chosen. The first and second landmark encounter in this base trial is at time *t* = 2s and *t* = 4.6s respectively. At time *t* = 1s (left), *t* = 4s (middle) and *t* = 7s (right) a multiplicative perturbation of size 50 % was introduced at the hidden layer.
- *Figure 3g*: Distribution 1
- *Figure 3i*: Landmarks and trajectories were sampled with the same parameters as distribution 1, except that the duration of test trials was extended from 10 s (100 timesteps) to 50 s (500 timesteps). In addition we only displayed trials with low error after the second landmark encounter. Low error trials were trials with maximum network localization error smaller than 0.5 rad, measured in a time window between 5 timesteps after the second landmark encounter until the end of the trial. In the figure, only the state-space trajectory after the second landmark encounter is displayed.
- *Figure 4a*: Distribution 2. Tuning curves were calculated using 20 bins of location/displacements and normalized individually for each neuron. The first time step in each trial and time steps with non-zero landmark input were excluded from the analysis.
- *Figure 4b*: Distribution 2. To determine the activity distribution, location/displacement in each condition was binned in 100 column bins and the response of each neuron was binned in 10 row bins. The resulting two-dimensional histogram was normalized to have equal sum for each column.
- *Figure 4c, top*: Performance was evaluated on 1000 trials from the distribution 2. For location, the decoder corresponded to the network location estimate. For displacement, the linear decoder was trained on 4000 separate trials from the same distribution.
- *Figure 4c, bottom*: Distribution 1 with 4000 trials to train the linear decoder and 1000 trials to evaluate it.
- *Figure 4d*: Location tuning curves were determined after the second landmark encounter using 1000 trials from distribution 2 and using 20 location bins. Tuning height specifies the difference between the tuning curve maximum and minimum, and tuning width denotes the fraction of the tuning curve with higher activation than the arithmetic mean of maximum and minimum.
- *Figure 4e, top*: Location tuning curves were determined after the second landmark encounter using 1000 trials from distribution 2 and using 20 location bins. Tuning curves were calculated separately for timesteps with clockwise and counterclockwise direction of movement.
- *Figure 4e, middle*: Location tuning curves were determined after the second landmark encounter using 1000 trials from distribution 2 and using 20 location bins. Velocity (irrespective of direction) was separated in 3 equal bins, corresponding to high, middle and low velocity. Tuning curves were calculated separately for each velocity bin.
- *Figure 4e, bottom*: Location tuning curves were determined for the whole trial using 1000 trials from distribution 1 and using 20 location bins. Uncertainty was measured by the posterior circular variance of the enhanced particle filter and binned in three equal bins. Tuning curves were calculated separately for each uncertainty bin.
- *Figure 4f*: This analysis was performed by evaluating the network of the external map task on the trajectory and environment distribution 1 of the internal map task. First, location tuning curves were determined after the second landmark encounter using 5000 trials from distribution 1 and using 50 location bins. Tuning curves were calculated separately for each of the four environment of the internal map task. Preferred location was determined to be the location corresponding to the tuning curve maximum. The density of preferred locations smaller than distance *d_min_* away from a landmark was then compared to to the density of preferred locations further away from landmarks.
- *Figure 4g*: Location tuning curves were determined after the second landmark encounter using 5000 trials from distribution 1 and using 20 location bins. The resulting tuning curves were shifted to have minimum value 0 and normalized to sum to one. The location entropy of each neuron was defined to be the entropy of the normalized location tuning curve. Neurons were split in two equal sets according to their location entropy, were neurons with low entropy were defined as “place cells” (PCs) and neurons with high entropy were defined as “non-place cells” (UCs). Between and across PCs and UCs absolute connection strength was calculated as the absolute value of the recurrent weight between non-identical pairs.
- *Figure 4h*: Location tuning curves were determined after the second landmark encounter using 5000 trials from distribution 1 and using 100 location bins. Preferred location was determined to be the location corresponding to the tuning curve maximum and neurons were sorted according to their preferred location. Shown is the hidden layer activation in an example trial with random trajectory and two landmark encounters, where hidden neurons are sorted according to their preferred location.
- *Figure 4i*: Location tuning curves were determined after the second landmark encounter using 5000 trials from distribution 1 and using 100 location bins. Preferred location was determined to be the location corresponding to the tuning curve maximum. Recurrent coupling of modes was defined by 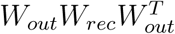, where *W_rec_* are the recurrent weights and *W_out_* are the output weights.
- *Figure S1a-g*: Analogous to Figure 4a-g, but for internal map network
- *Figure S1c, bottom*: A multinomial regression decoder was trained on 4000 trials from distribution 1 (the training distribution of the internal map task) to predict from hidden layer activities which of the four possible environments was present. The performance was evaluated on separate 1000 test trials sampled from the training distribution. The graph shown shows the performance conditioned on the number of landmark visits.
- *Figure S2a-g*: Analogous to Figure 4a-e, 4g and 3g respectively, but for non-negative network.
- *Figure S3a-f*: Analogous to Figure 3a-d and 3f-g respectively, but for internal map network.
- *Figure S4a*: Distribution of non-diagonal recurrent weights for randomly initialized (untrained), external map, internal map and non-negative network. The *k*-value measures denotes excess kurtosis, a measure of deviation from Gaussianity (*k* = 0 for Gaussian distributions).
- *Figure S4b*: Scatterplot of real and imaginary part of complex eigenvalues of recurrent weight matrix for randomly initialized (untrained), external map, internal map and non-negative network.
- *Figure S5a:* Normalized location tuning curves after the second landmark encounter for all 128 hidden neurons for the external map network.
- *Figure S5b*: Normalized signed velocity tuning curves after the second landmark encounter for all 128 hidden neurons for the external map network.
- *Figure S6*: 5000 trials from the hidden activity of a recurrent network with random initial hidden state, frozen large random recurrent weights and without inputs. The recurrent weights were sampled i.i.d. from a uniform distribution *W_h,ij_*~*U* ([-1,1]), then fixed across trials. The initial hidden state across trials was sampled from from a uniform distribution *h_t=0,i_*~*U* ([-1,1]).

**Figure S1:**
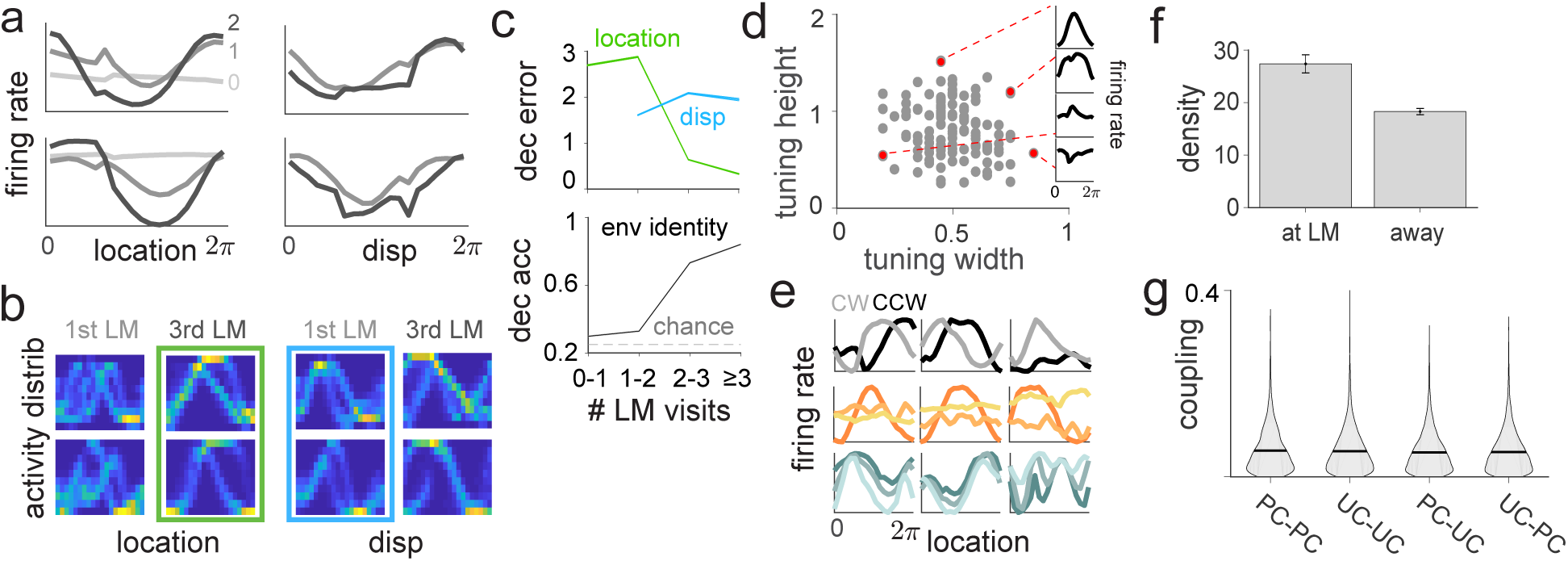
Hidden layer tuning curves for internal map network **a - f**, Analogous to Figure 4a-e and 4g

**Figure S2:**
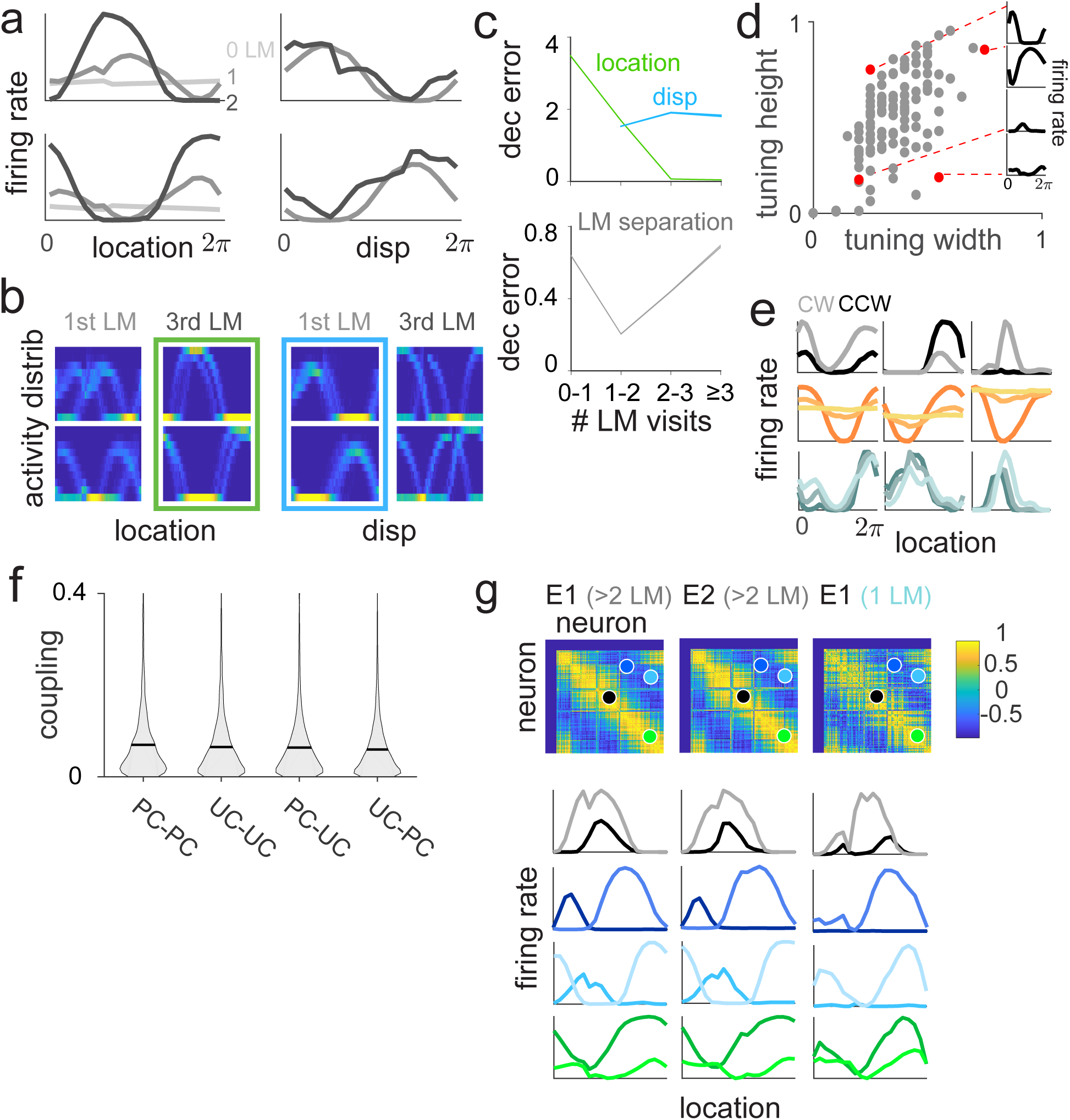
Hidden layer tuning curves for non-negative network **a - g**, Analogous to Figure 4a-e and 4g

**Figure S3:**
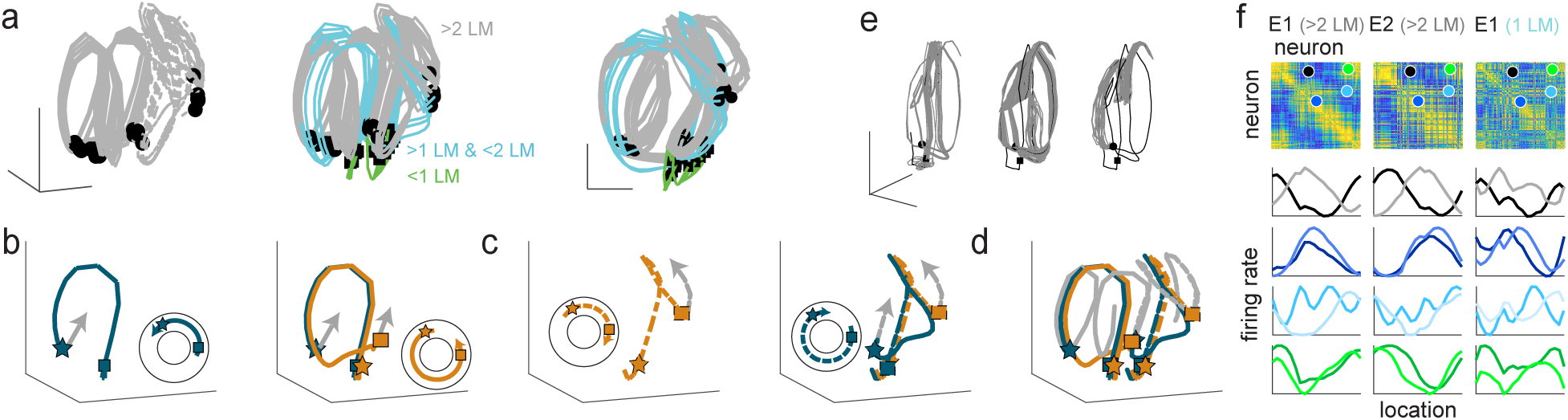
Population dynamics in hidden layer for internal map network **a - f**, Analogous to Figure 3a-d and 3f-g, respectively

**Figure S4:**
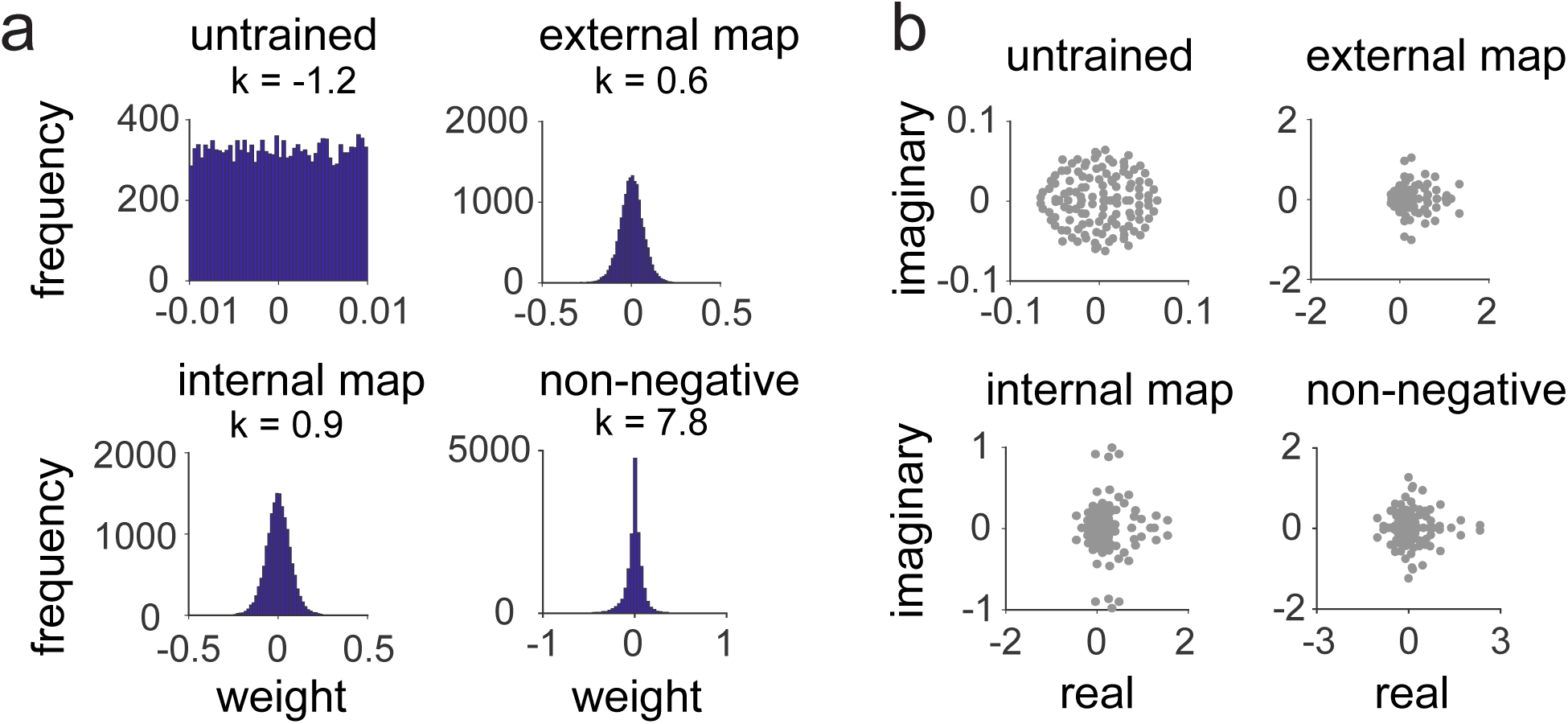
Recurrent weight distribution.

**Figure S5:**
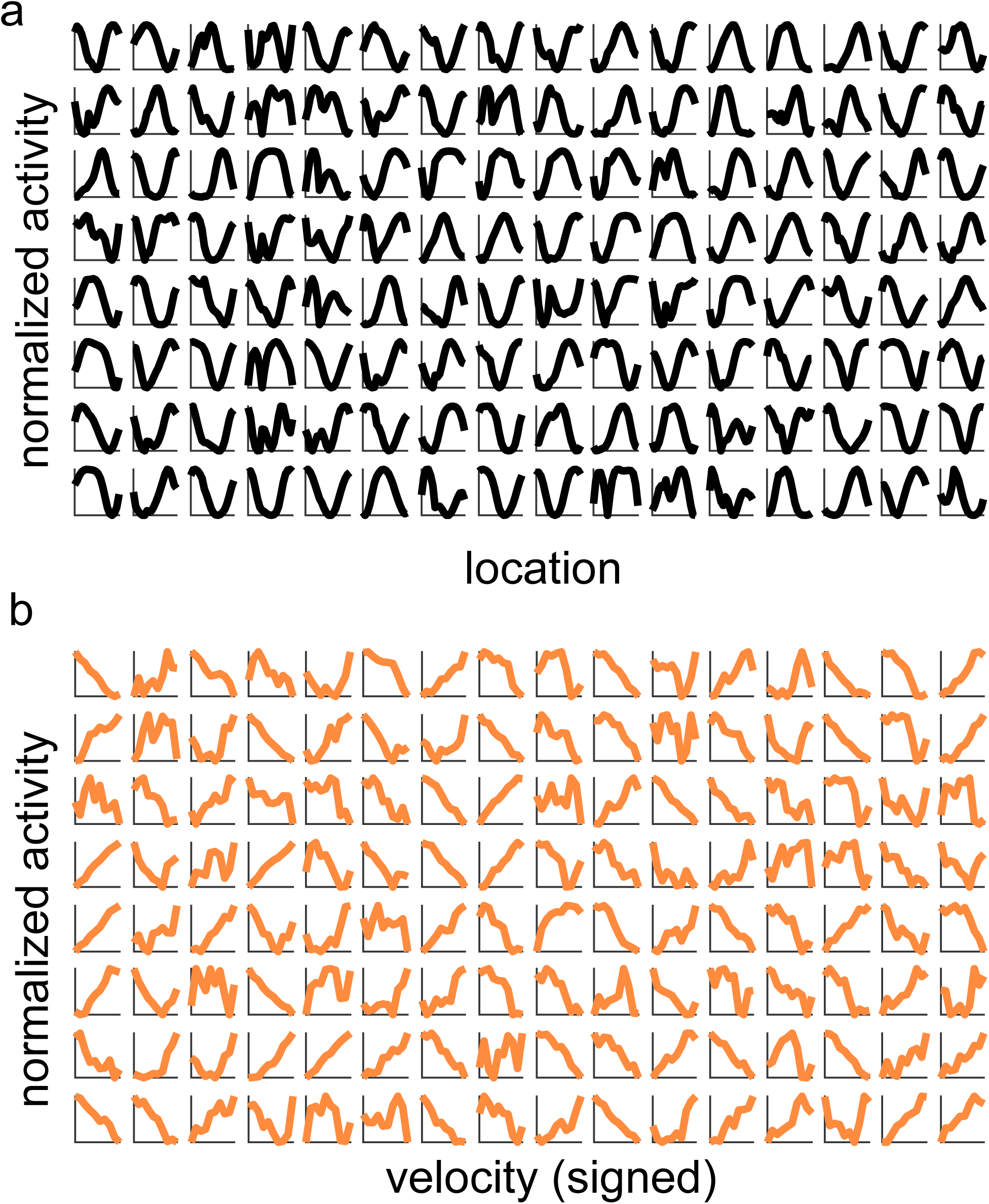
Full set of tuning curves for all 128 hidden neurons. **a**, Location tuning curves, **b**, Velocity tuning curves (sign of velocity corresponds to direction)

**Figure S6:**
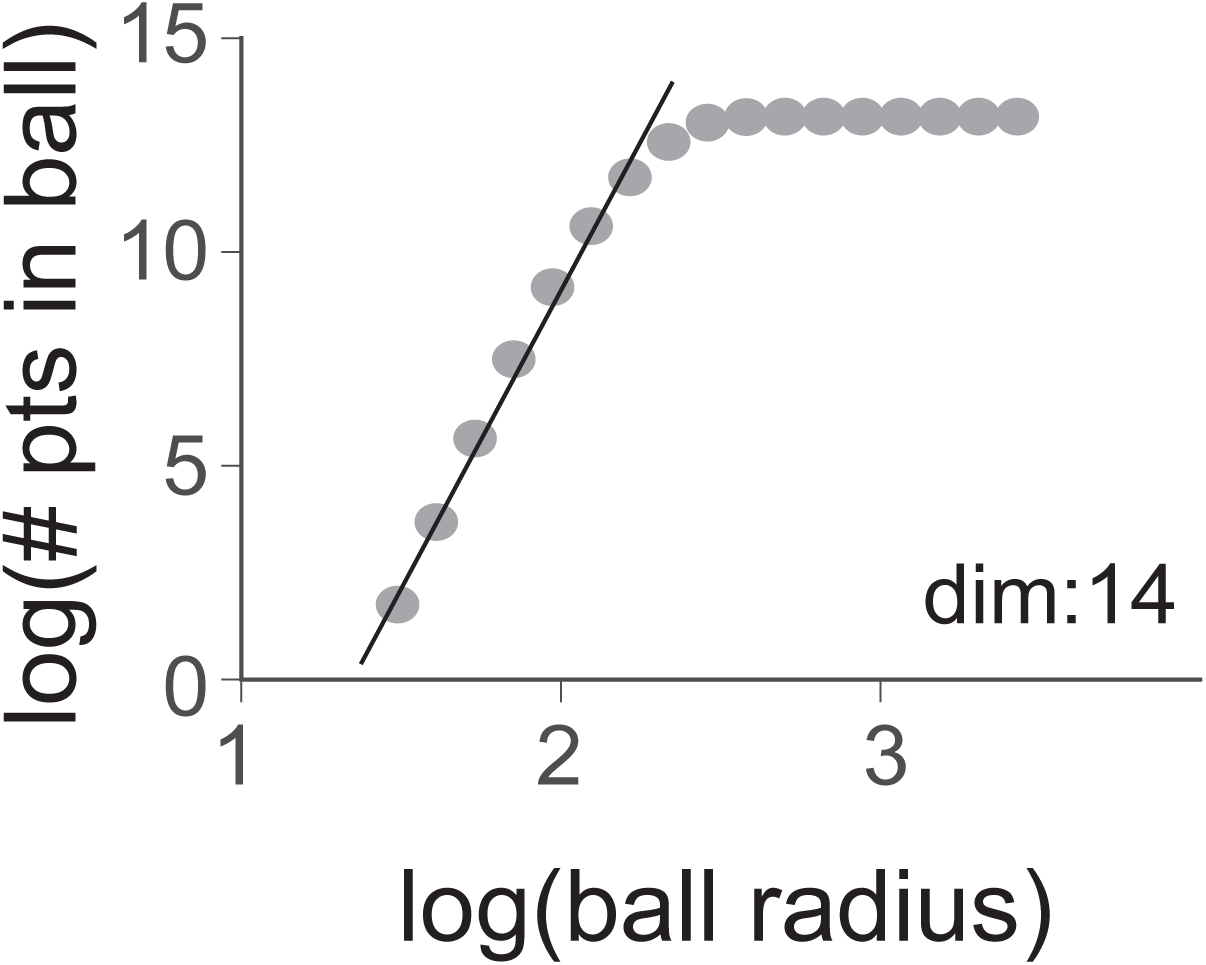
Correlation dimension for network with large random weights and random initialization.

